# Time-resolved coupling between connectome harmonics and subjective experience under the psychedelic DMT

**DOI:** 10.1101/2024.05.30.596410

**Authors:** Jakub Vohryzek, Andrea I. Luppi, Selen Atasoy, Gustavo Deco, Robin L. Carhart-Harris, Christopher Timmermann, Morten L. Kringelbach

## Abstract

Exploring the intricate relationship between brain’s structure and function, and how this affects subjective experience is a fundamental pursuit in neuroscience. Psychedelic substances offer a unique insight into the influences of specific neurotransmitter systems on perception, cognition and consciousness. Specifically, their impact on brain function propagates across the structural connectome — a network of white matter pathways linking different regions. To comprehensively grasp the effects of psychedelic compounds on brain function, we used a theoretically rigorous framework known as connectome harmonic decomposition. This framework provides a robust method to characterize how brain function intricately depends on the organized network structure of the human connectome. We show that the connectome harmonic repertoire under DMT is reshaped in line with other reported psychedelic compounds - psilocybin, LSD and ketamine. Furthermore, we show that the repertoire entropy of connectome harmonics increases under DMT, as with those other psychedelics. Importantly, we demonstrate for the first time that measures of energy spectrum difference and repertoire entropy of connectome harmonics indexes the intensity of subjective experience of the participants in a time-resolved manner reflecting close coupling between connectome harmonics and subjective experience.

## Introduction

Understanding how subjective experience arises from the dynamic interplay of brain structure and function is a central question in neuroscience. In combination with non-invasive neuroimaging, psychedelic substances offer a powerful window to interrogate how specific neurotransmitter systems shape brain function to influence perception, cognition, and consciousness. Crucially, the changes in brain function exerted by neurotransmitter engagement propagate throughout the brain according to the network of white matter pathways between regions: the human structural connectome. Therefore, understanding the effects of psychedelic compounds on brain function involves bridging structure and function across multiple levels.

A theoretically rigorous way to characterise how brain function depends on the underlying network organisation of the human connectome is provided by the framework of connectome harmonic decomposition (CHD). Mathematically, CHD represents functional signals in terms of their dependence on the underlying structural connectome. In other words, CHD is a change of basis functions, analogous to the Fourier transform that transforms a signal from the time domain into the domain of temporal frequencies. Likewise, CHD transforms brain signals from the spatial domain, into the domain of connectome frequencies. CHD explicitly expresses brain activity in terms of multi-frequency contributions from the underlying structural network: each connectome harmonic is a distributed activation pattern characterized by a specific spatial scale (frequency). Low-frequency (coarse-grained) connectome harmonics indicate that the functional signal reflects global connectivity patterns in the underlying structural connectome. In turn, high-frequency (fine-grained) connectome harmonics indicate a divergence between the spatial organisation of the functional signal coupled to the (coarse-grained) underlying network structure: nodes may exhibit different functional signals even if they are closely connected to the same structural network.^1^

Recent work has consistently demonstrated two prominent effects of psychedelics on the connectome harmonic landscape of the human brain. First, the serotonergic psychedelics, LSD and psilocybin, as well as the atypical psychedelic, ketamine, consistently induce a reduction in the contribution of low-frequency (large-scale) harmonics, and a corresponding increase in the contribution of high-frequency (fine-grained) harmonics.^2–4^ This evidence is also in line with additional reports of LSD-induced structure-function decoupling^5^ where others have interpreted a shift away from low-frequency harmonics in favour of high-frequency ones as decoupling of brain activity from the underlying structural connectivity^1, 4^; or at least the major white tractography. Second, psychedelics induce a broadening of the repertoire of connectome harmonics that contribute to spontaneous brain activity^4^ alongside evidence of increases in the spatio-temporal metastability of brain function in the psychedelic state.^6, 7^

Here, we hypothesise that as a potent serotonergic psychedelic, DMT will reshape the connectome harmonics in line with the effects previously reported for LSD and psilocybin, as well as the atypical psychedelic, ketamine. Namely, we predict a decreased contribution from low-frequency harmonics under the effects of DMT, and instead an increase in the contribution of high-frequency harmonics. We also hypothesise that like other psychedelics, DMT will increase the diversity (entropy) of the repertoire of connectome harmonics.

A crucial feature of the effects of intravenous (IV) DMT, that makes it especially valuable for scientific investigation is that, whereas oral LSD- and psilocybin-induced effects have a slow onset and can last for several hours, the effects of IV DMT are relatively more contracted and temporally predictable. IV DMT has a fast onset and reliably short duration of approximately 8 minutes for the dosage and injection parameters used here. This feature of DMT makes it possible to obtain dynamic ratings of the intensity of subjective experience over time, and then relate these data to the corresponding time-resolved changes in connectome harmonics - since CHD analysis is also applicable on a dynamic timepoint-by-timepoint basis. Recent results have shown that neural changes in connectome harmonic signature reflect changes in subjective experience.^4^ However, those results were time-averaged across the entire scan duration. Therefore, here we capitalise on the unique temporal resolution offered by DMT to test a stronger hypothesis: that the neural changes in connectome harmonic composition - as described by energy spectrum and repertoire entropy - will be related to behavioural changes in intensity ratings, not just on average, but rather in a dynamic timepoint-by-timepoint manner, reflecting close coupling between connectome harmonics and subjective experience.

## Methods

### DMT dataset

The complete description of the participants, experimental design and acquisitions parameters can be found in.^8^ Below, we provide a succinct account of consistent information.

### Participants

A group of 25 participants was recruited in a single-blind, placebo-controlled and counter-balanced design. Subjects were considered for the study unless they were younger than 18 years of age, lacked experience with a psychedelic, had a previous negative response to a psychedelic and/or currently suffered from or had a history of psychiatric or physical illness. Out of the 25 participants, 20 completed the whole study (7 female, mean age = 33.5 years, SD = 7.9). A further 3 subjects were excluded due to excessive motion during the 8 minutes DMT recording (more than 15% of volumes scrubbed with framewise displacement (FD) of 0.4 mm).

### Experimental Paradigm

In total, all subjects were scanned on two days, two weeks apart, each consisting of two scanning sessions. The initial scan lasted 28 minutes with the 8th minute marking the intravenous administration of either DMT or placebo (saline) (50/50 DMT/placebo), single bolus lasting 60 seconds. Subjects were asked to lay in the scanner with their eyes closed (wearing an eye-mask). After the recording, assessment of subjective effects was carried out. The second session was identical to the first except for the assessment of subjective intensity scores at every minute of the recording. The experimental design also included simultaneous EEG recording during the sessions (see Figure 1).

**Fig 1.**
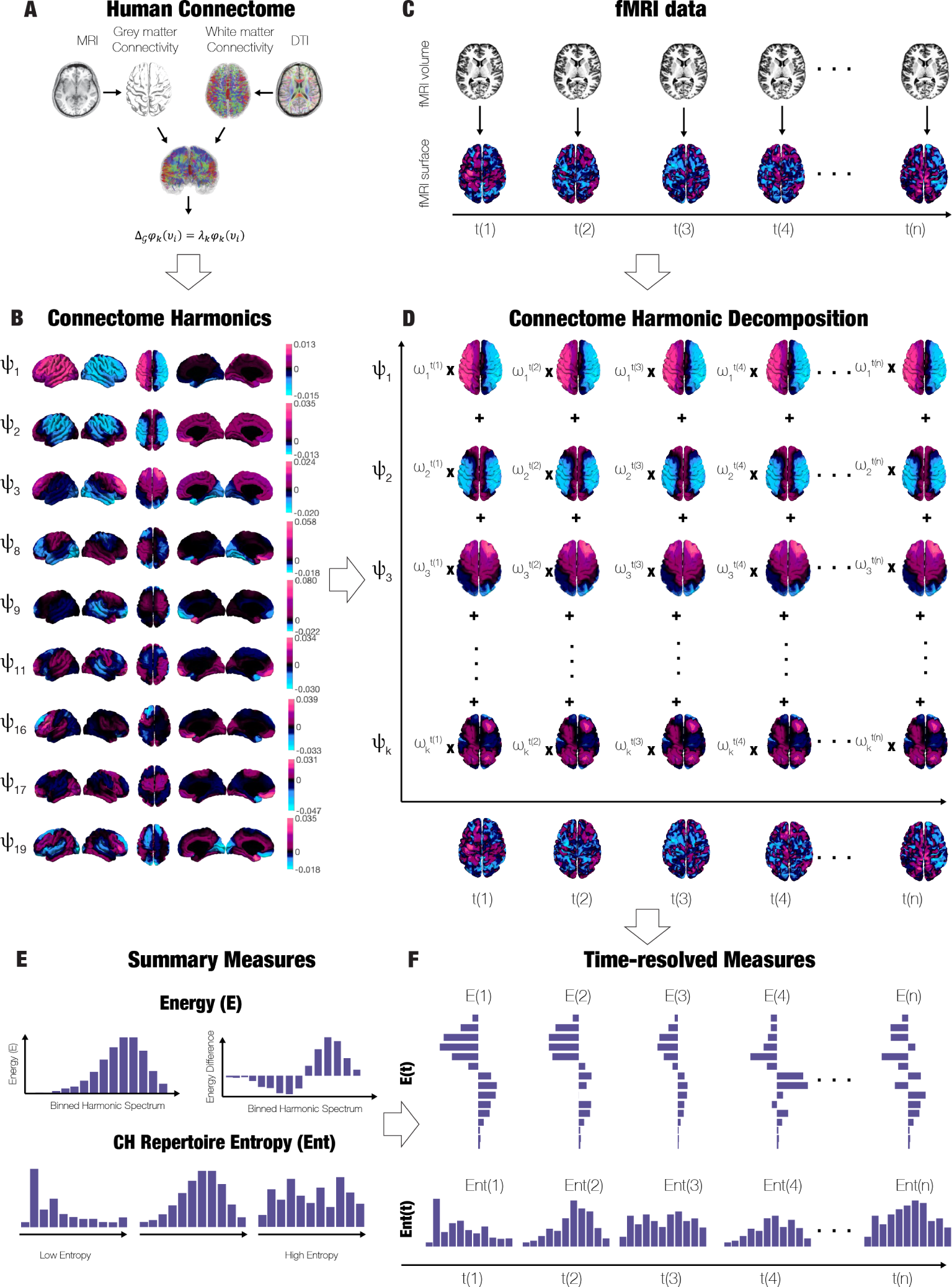
Study Overview: **A)** Human Connectome constructed from structural and diffusion MRI. **B)** Connectome Harmonics computed from the eigendecomposition of Laplacian operator applied to the human connectome. **C)** Functional MRI projected from MNI volumetric space to Freesurfer surface space. **D)** Connectome Harmonic Decomposition summarising fMRI timeseries as a linear summation of individual harmonics and their weights. **E)** Summary measures for interpreting the Connectome Harmonic Decomposition. Namely, the energy spectrum, energy spectrum difference and CH repertoire entropy **F)** Time-resolved measures applied to the entire course of the fMRI recordings.

### fMRI Acquisition Parameters

The experiment was performed on a 3T scanner (Siemens Magnetom Verio syngo MR 12) with compatibility for EEG recording. A T2^∗^-weighted echo planar sequence was used. In brief, the parameters were as follows: TR/TE = 2000ms/30ms, acquisition time = 28.06 minutes, flip angle = 80°, voxel size = 3×3×3 mm^3^ and 35 slices with 0 mm interslice distance. T1-weighted structural scans of the brain were also acquired.

### FMRI Pre-processing

For fMRI pre-processing, a pipeline previously developed for an LSD experiment was used, which can be accessed in the supplementary information of.^9^ Briefly, the following steps were applied 1) despiking, 2) slice-timing correction, 3) motion correction, 4) brain extraction, 5) rigid body registration to structural scans, 6) non-linear registration to 2mm MNI brain, 7) motion-correction scrubbing, 8) spatial-smoothing (FWHM) of 6 mm, 9) band-pass filtering into the frequency range 0.01-0.08 Hz, 10) linear and quadratic detrending, 11) regression of 9 nuisance regressors (3 translations, 3 rotations and 3 anatomical signals).

### Psilocybin Dataset

A complete description of the psilocybin study protocol can be found in the original paper.^10^ In brief, nine participants were considered for this analysis following rigorous exclusion criteria; no younger than 21 year of age, pregnancy, history of psychiatric disorders, cardiovascular disease, substance dependence, claustrophobia, blood or needle phobia or adverse response to psychedelics. Furthermore, participants were excluded if their mean framewise displacement (FD) exceeded 0.4 mm. Two eye-closed scans, separated by seven days, were performed for each participant. In a counterbalanced design each participant was given intravenously either psilocybin (2 mg dissolved in 10 ml saline, 60 s) or saline (10 ml saline, 60 s). The recording session lasted 12 minutes with the 6th minute marking the solution infusion. For the study last five minutes (post the 60 s infusion) were analysed and a control pre-injection recording was matched accordingly to be 5 minutes as done in the original study.^10^ We analysed segments of 100 timepoints pre- and post-psilocybin injection (TR = 3 seconds).

### LSD Dataset

A complete description of the LSD study protocol can be found in the original paper.^11^ Briefly, fifteen subjects were considered for this analysis with key exclusion criteria; no younger than 21 year of age, pregnancy, history of psychiatric disorders, cardiovascular disease, substance dependence, claustrophobia, blood or needle phobia or adverse response to psychedelics, previous experience with serotonergic psychedelics or use within 6 weeks of first scanning. Furthermore participants were excluded if their mean framewise displacement (FD) exceeded 0.4 mm. In a counter-balanced design each participant was given intravenously either LSD (75 *µ* g in 10 ml saline, 120 s) or saline (10 ml saline, 120 s). After 60 minutes acclimatisation in the scanner, there were three resting-state recording sessions; one with music interleaved in-between two standard resting-state sessions. Only results of the first resting-state recording session are considered here as done in the original study.^9^ The scan lasted 7.23 minutes with a TR of 2 seconds, resulting in 217 timepoints pre- and post-LSD injection.

### Structural Connectome Construction

For the construction of group connectome harmonics, an independent cohort of 10 participants (6 female, 22-35 years) was used from the Human Connectome Project, WU-Minn Consortium (Principal Investigators: David Van Essen and Kamil Ugurbil: 1U54MH091657). This project was made possible by funding from the sixteen NIH Institutes and Centres supporting the NIH Blueprint for Neuroscience Research; and by the McDonell Centre for Systems Neuroscience at Washington University. Both structural and Diffusion Tractography Imaging (DTI) data was used for the construction of connectomes with pre-processing according to the minimal pre-processing guidelines of the HCP protocol.^12^

For the estimation of the connectome harmonics, we used the identical workflow as in Atasoy et al. (2017).^2^ In general, this consisted of combining local, surface based, and long-range white-matter connectivity in a sparse vertex-based representation. In brief, cortical surface reconstruction from high-resolution T1-weighted MRI of individual participants was carried out with Freesurfer software. Then, each participant’s cortical surface was registered to the 1000-subject group template yielding a common-space mesh of 10,242 vertices in each hemisphere. For the white-matter cortico-cortical fibers, deterministic tractography was applied to the DTI data of individual subjects (resolution 1.25 mm) with Matlab implementation of Vista Lab, Stanford University. For the tractography itself, eight seeds were initialised in each vertex (total of 20,484) with the termination criteria being either fractional anisotropy (FA) below 0.3, minimum track length of 20 mm and a maximum angle of 30° between two adjacent tracking steps.

### High-resolution alternative reconstruction of the human connectome

To demonstrate that our results are not fundamentally dependent on this specific operationalisation of the human connectome, we also used an alternative representative human connectome, following the same procedures as in our recent report.^4^ The alternative connectome was constructed from multi-shell diffusion-weighted imaging data from 985 subjects of the HCP 1200 data release (http://www.humanconnectome.org/), each scanned for approximately 59 minutes. This represents a nearly 100-fold increase in sample size compared with the original connectome used for connectome harmonic decomposition.^2, 13^ We refer to the human connectome constructed from these data as the HCP-985 connectome. Acquisition parameters are described in detail in the relative documentation (http://www.humanconnectome.org/), and the dMRI data were preprocessed and made available as part of the freely available Lead-DBS software package (http://www.lead-dbs.org/). For the reconstruction of long-range white matter tracts of each individual, we followed the procedures previously used on these data:^4^ the diffusion data were processed using a generalized q-sampling imaging algorithm implemented in DSI Studio (http://dsi-studio.labsolver.org). A white-matter mask was obtained from segmentation of the T2-weighted anatomical images, which were co-registered to the b0 image of the diffusion data using SPM12. In each HCP participant, 200,000 fibres were sampled within the white-matter mask, using a tracking method that previously achieved the highest valid connection score among 96 methods submitted from 20 different research groups in a recent open competition.^14^ Finally, the fibres were transformed into standard Montreal Neurological Institute (MNI-152I) space using Lead-DBS software.^15^ The remaining procedures for obtaining individual connectomes and aggregating them into a group-average representative connectome, and subsequent connectome harmonic decomposition, were the same as described above.

### Randomised connectome

To demonstrate the importance of the specific topology of the human connectome, obtained by combining local grey matter connectivity and long-range white matter fibres, we also tested whether our results would replicate when using harmonics obtained from a randomised connectome.^4^ Before performing Laplacian decomposition (as described in the next section), the original connectome was therefore turned into a random network using the degree-preserving procedure implemented in the Brain Connectivity Toolbox.^16^ Harmonics were then extracted from Laplacian eigendecomposition, and the full connectome harmonic decomposition pipeline was followed.

### Derivation of Connectome Harmonics

Connectome Harmonics were computed from the structural connectome described as a graph, ℜ = (*ν, ε*), of vertices *ν* = {*v_i_*| ∈ 1, …, *n*} and edges *ε* = {*e_ij_*| ∈ *ν* × *ν*}. The graph’s edges represent 1) local connectivity, defined by 6 nearest neighbours of each vertex on the cortical surface, and 2) long-range connectivity as determined by tractography in terms of cortico-cortical fibres. The graph’s edges were further binarised and symmetrised and represented in an unweighted and undirected adjacency matrix, *A*, as follows:

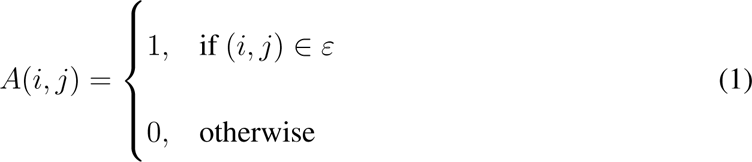

where *i*,*j* are the indices of the adjacency matrix *A* (20,424 × 20,424). In order to obtain a group adjacency matrix 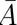, the individual adjacency matrices of the 10 subjects were averaged. Furthermore, the discrete counterpart of the Laplace operator Δ, applied to the structural connectivity (local and long-range) of the human brain 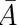, is estimated by computing the symmetric Laplacian 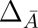 in the following manner:

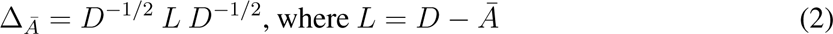

where the *D* is the diagonal degree matrix, 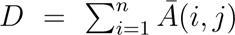. Lastly, the connectome harmonics, *ψ_k_*, *k* ∈ 1, …, *n* were defined and computed as the eigenvectors of the following eigen-value problem,

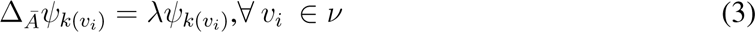

where *λ, k* ∈ 1, …, *n* are the associated eigenvalues of 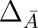.

### Connectome Harmonic Decomposition of Resting-state fMRI

The derived connectome harmonics can be used to represent resting-state fMRI in each of the DMT conditions. First, resting-state fMRI is projected from its MNI voxel-space to the Freesurfer surface vertex-space using the HCP command *-volume-to-surface-mapping*. The resulting timecourse can be represented as *F*(*v, t*) for every vertex *v* ∈ *ν*. Once in the same space, the fMRI activity *F*(*v, t*) at every time step *t* ∈ [1, …, *T*] can be described as a sum of weighted contributions *α_k_* of individual connectome harmonics *k*. This fMRI decomposition can be described in the following format,

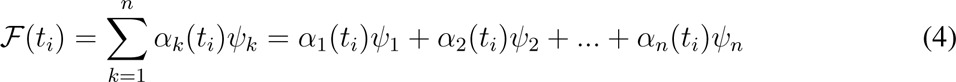

with *α_k_* being the contribution of *k* connectome harmonic *ψ_k_* to the fMRI activity *F*(*t_i_*) at time *t_i_*. Formally, the connectome harmonic contributions are described as *α_k_*(*t*) = ⟨*F*(*t*), ψ*_k_*⟩.

### Measures

Further analysis describes several measures which summarise different aspects of the decomposed fMRI timecourses in terms of the contributions of the connectome harmonic spectrum.

### Power and Energy

The connectome harmonic weight *α_k_* at each time *t* represents the strength of a given connectome harmonic *ψ_k_* of that particular fMRI pattern F(*v, t*) at the same time *t* and its absolute value can be defined as power, *P* (*ψ_t_, t*) = |*α_k_*(*t*)|. To further estimate the connectome harmonic contribution *α_k_* in relation to its eigenvalue *λ_k_*, a measure of energy, *E*, is defined as the square of connectome harmonic contribution and its intrinsic energy,(*λ*), in the following way 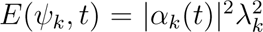. Note that both Power and energy are computed for every connectome *ψ_k_* at every timepoint *t* across the fMRI timecourse *F*(*v, t*). Moreover,we used the energy spectrum averaged over 15 bins of varying logarithmic scale for the 18k connectome harmonics. Subsequently, the energy spectrum is defined as the difference between the 8 minutes recordings of the pre- and post-recording sessions for the DMT and placebo groups.

To ensure that our results are not specific to the chosen number of bins, we also replicate our main analysis using 25 bins instead of 15. Additionally, to ensure that our results were not unduly influenced by potential aliasing effects introduced by the use of high-resolution diffusion data, for the HCP-985 analyses we only used the first 14 logarithmically spaced bins (instead of 15 as for the previous analyses),^4^ showing that our results are not critically dependent on the precise number of bins. Likewise, for the analysis with randomised connectome we also only used the first 14 logarithmically spaced bins.

### Repertoire Entropy of Connectome Harmonics

The repertoire entropy of CH, *H*, was defined as the time-averaged histogram entropy of the binned (15 logarithmic bins) power distribution, *P*, at each condition i.e. 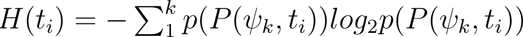. The bin choice for both measures was chosen historically as a common frame of reference for comparison with previous literature.^3^ Intuitively, the flatter the distribution the higher the repertoire entropy of connectome harmonics, while the more concentrated distribution to a given frequency range the lower the repertoire entropy of connectome harmonics.

### Data-driven extraction of multivariate connectome harmonic signatures

Partial least squares (PLS) is a multivariate statistical analysis used to identify relationships between one or more targets (Y) and a set of predictor variables X. This method extracts principal components as linear combinations of variables in each set that maximally covary with each other. In the present case, for both DMT and placebo, X was the matrix of 15 binned energy values (see above) for each individual (averaged over timepoints), and Y was the vector of binary classification between the two states (DMT or placebo) – making this an application of Partial Least Squares Discriminant Analysis (PLS-DA), owing to the binary nature of Y.^17^ The first principal component extracted by PLS-DA represents the single most discriminative pattern present in the data, in terms of distinguishing observations (subjects) belonging to the two different classes (placebo versus DMT).

## Results

Using connectome harmonics as the spatial basis of brain activity, it is possible to describe the temporal evolution of connectome harmonics in terms of their contribution. Here, we use CHD to describe the spatio-temporal changes of the DMT-induced state in terms of its connectome harmonic spectrum and repertoire diversity (entropy).

### The DMT-induced State Suppresses Low-level Harmonics and Increases High-level Harmonics

We first estimated the connectome harmonic energy spectrum of each condition (DMT pre/post and PCB pre/post) across all timepoints and subjects. Following the established procedure for connectome harmonic analysis,^2–4^ we then binned the connectome harmonic spectrum into 15 logarithmically spaced bins and obtained the harmonic profiles.

For the DMT-induced state, a range of low frequency harmonic bins (*k* ∈ [1,…,10^2^]) were found significantly suppressed as opposed to the pre-DMT condition (p-value < 0.05, Bonferroni corrected paired t-test). No significant differences were observed in the placebo condition. A mirror opposite change was observed in the high frequency harmonic bins, whereby a range of *k* ∈ [10^3^,…,10^4^] was found significantly increased (p-value < 0.05, Bonferroni corrected paired t-test). Again, no significant differences were observed for the placebo condition (Figure 2, A). This profile change across quantized harmonic bins can be further explored by looking at the energy differences across the DMT conditions of each subject while comparing it to the placebo condition difference. Remarkably, a similar suppression of the lower-harmonics *k* ∈ [1,…,10^2^] (p-value < 0.05, Bonferroni corrected paired t-test) and increase of the higher harmonics *k* ∈ [10^3^,…,10^4^] (p-value < 0.05, Bonferroni corrected paired t-test) are observed with non-significant results for bins at the inflexion point [10^2^,…,10^3^] (Figure 2, B). Furthermore, these energy distribution changes of CH under DMT are robust to the choice of connectome as the results are consistent with the analysis performed with the 985 HCP participant connectome (SI Figure S1). Lastly, these energy distribution changes are also consistent when comparing the DMT and placebo post-injection conditions for both the original and 985 HCP participant connectomes (SI Figure S2).

**Fig 2.**
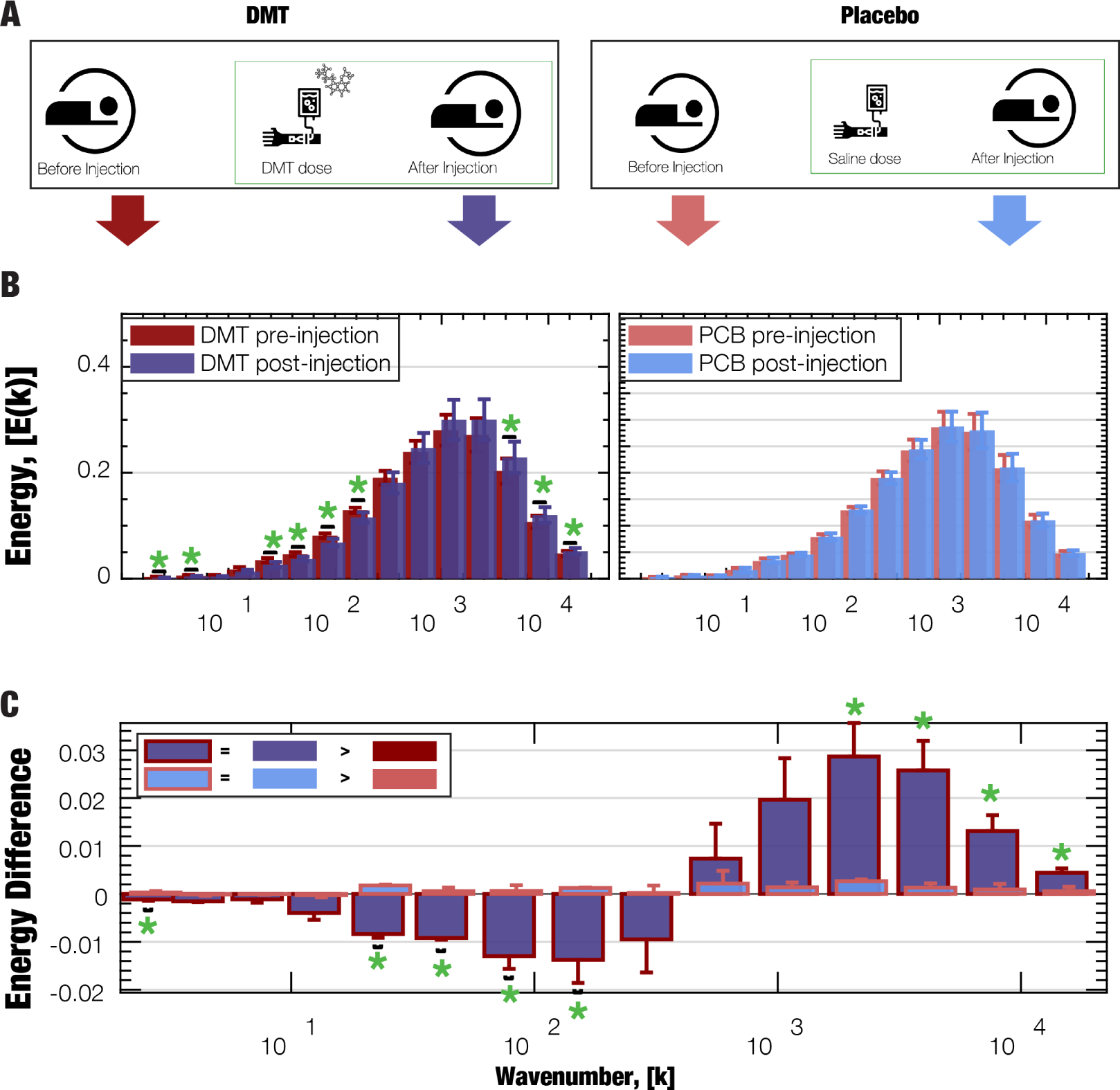
DMT-induced Energy Changes in the Connectome Harmonic Spectrum: **A)** A decrease of energy for low-frequency harmonics *k* ∈ [1,…,10^2^] (p-value < 0.01, Bonferroni corrected paired t-test) and an increase of energy for high-frequency harmonics *k* ∈ [10^3^,…,10^4^] (p-value < 0.01, Bonferroni corrected paired t-test) were observed. No significant changes was observed in the placebo conditions. **B)** Furthermore, energy differences between the DMT and placebo conditions were observed with decreases in low-frequency harmonics *k* ∈ [1,…,10^2^] (p-value < 0.01, Bonferroni corrected paired t-test) and increases in high-frequency harmonics *k* ∈ [10^3^,…,10^4^] (p-value < 0.01, Bonferroni corrected paired t-test). No significant differences were observed at the inflexion point [10^2^,…,10^3^]

### Contextualising DMT-induced Changes in Connectome Harmonic Spectrum Against Other States of Consciousness

DMT is a classical serotonergic psychedelic, pharmacologically related to psilocybin and LSD. Here, we show that the connectome harmonic signature of DMT coincides with the previously reported signatures of LSD and psilocybin.^3, 4^

The previous analyses consider each CH bin in isolation. However, it is clear that the overall pattern that emerges from considering all bins together is just as meaningful—if not more so. To take into account the full spectrum of connectome harmonic changes at the same time, we followed our previous work^4^ and turned to Partial Least Squares Discriminant Analysis (PLS-DA): this data-driven technique allowed us to extract the multivariate patterns of connectome harmonic energy that maximally distinguish between DMT and placebo (termed “multivariate signatures”, MVS). This approach previously revealed the existence of two mirror-reversed multivariate patterns characterising loss of consciousness (anaesthesia and disorders of consciousness) and the psychedelic state (LSD and the atypical psychedelic, ketamine).^4^ Here, we sought to test the hypothesis that, as a psychedelic, DMT would align positively with the multivariate signatures of LSD and psychedelic doses of ketamine, and negatively with the signatures of unconsciousness (awake vs propofol, and DOC fMRI+ vs fMRI-, corresponding to brain-injured patients who can versus cannot provide in-scanner evidence of responding to linguistic commands). We pursued this hypothesis by projecting each subject’s connectome harmonic energy spectrum onto a given MVS (thereby measuring the correspondence between them), and then comparing the value of this projection across DMT and placebo conditions. Supporting our hypothesis, we clearly found that the multivariate connectome harmonic signature that best distinguishes DMT from placebo, coincides with the analogous signatures of LSD and psychedelic ketamine. Conversely, the DMT signature (placebo vs DMT) is the opposite of the signatures obtained by comparing wakefulness against propofol anaesthesia, or fMRI-responsive versus unresponsive DOC patients (Figure 3). Furthermore, these results are reproduced when using the 985 HCP participants connectome as the structural basis (SI Figure S7)

**Fig 3.**
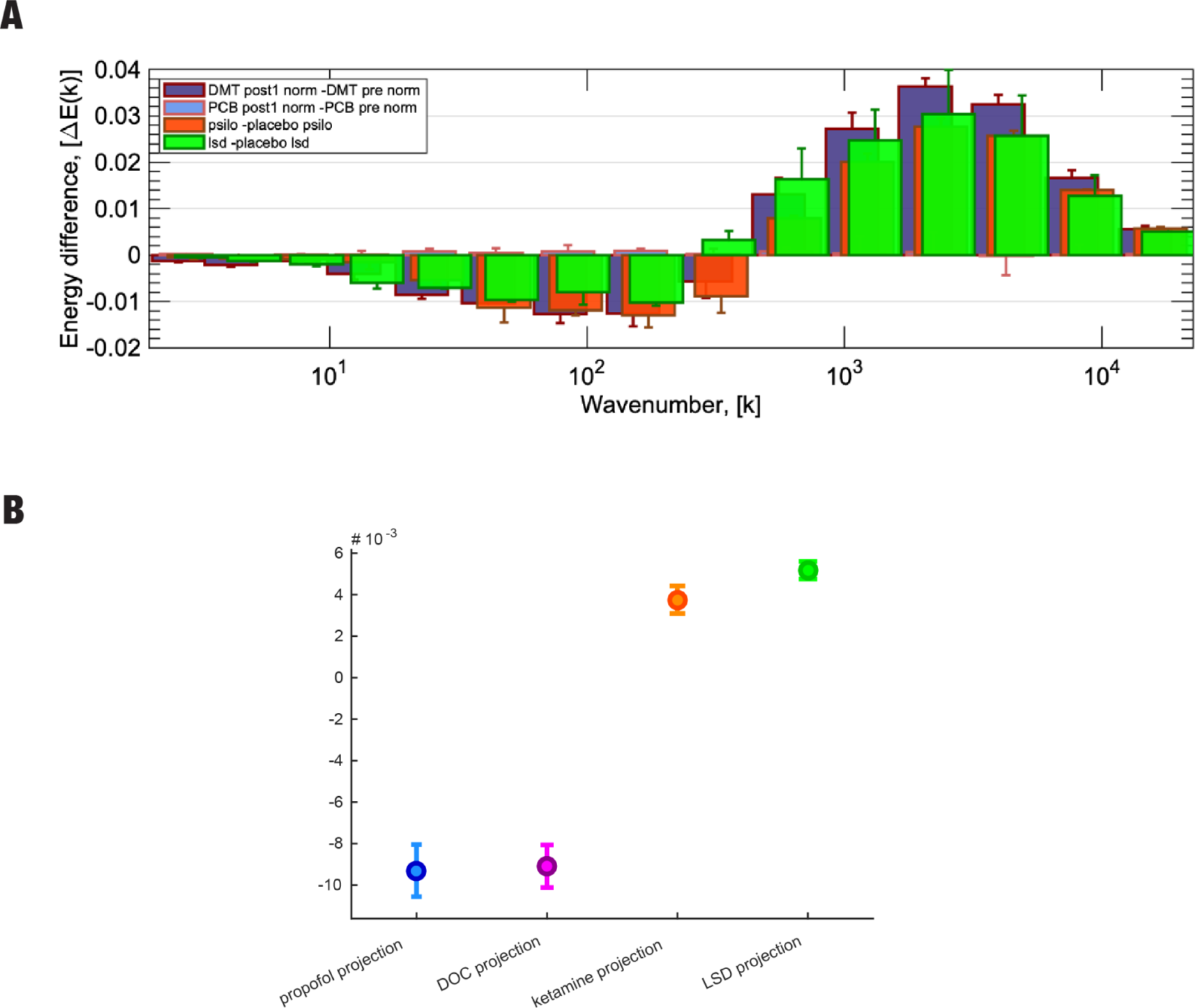
Contextualising the Connectome Harmonic signature of DMT with other altered brain states: **A)** The Connectome Harmonic signature of DMT (energy difference) is shown alongside corresponding signatures of psilocybin and LSD-induced states previously reported in^3^, to enable visual comparison. The control placebo condition from the DMT study is also shown, to demonstrate that effects are specific to altered states of consciousness. **B)** Fixed effects (and 95% CI) of projections (dot product) between the multivariate connectome harmonic signature of DMT, and four other states previously investigated by^4^ : anaesthesia (blue), DOC patients (violet), ketamine (orange), and LSD (green); all p < 0.001.

### DMT Enhances the Diversity of Connectome Harmonics Repertoire

The prominent entropic brain account of psychedelic action posits that psychedelics exert their subjective effects at least in part by increasing the diversity (entropy) of spontaneous brain activity and connectivity, which would then translate to greater richness of subjective experience^18, 19^ or ‘phenomenal consciousness’.^20^

Here, we therefore investigate whether DMT, a psychedelic, also increases the entropy of the connectome harmonic repertoire, as predicted by the entropic brain hypothesis and shown here in EEG data, where the entropy of spontaneous brain activity and subjective ‘richness’ were strongly correlated’.^21^ We computed the normalised CH repertoire entropy for each condition (pre/post DMT and pre/post placebo). CH repertoire entropy increased in the DMT-induced state compared to the other three conditions (Pre/Post DMT: p-value < 10^−4^, Pre PCB/Post DMT:-value < 10^−4^, Post PCB/Post DMT: p-value < 10^−4^, paired t-test). Furthermore, the result was strengthened by comparing the CH repertoire entropy difference between post/pre DMT and post/pre placebo where an increase was observed as well (Diff. in Pre-Post DMT and Pre-Post PCB: p-value < 10^−5^, paired t-test). Furthermore, the changes in CH repertoire entropy of CH under DMT are robust to the choice of connectome as the results are consistent with the analysis performed with the 985 HCP participant connectome (SI S3).

**Fig 4.**
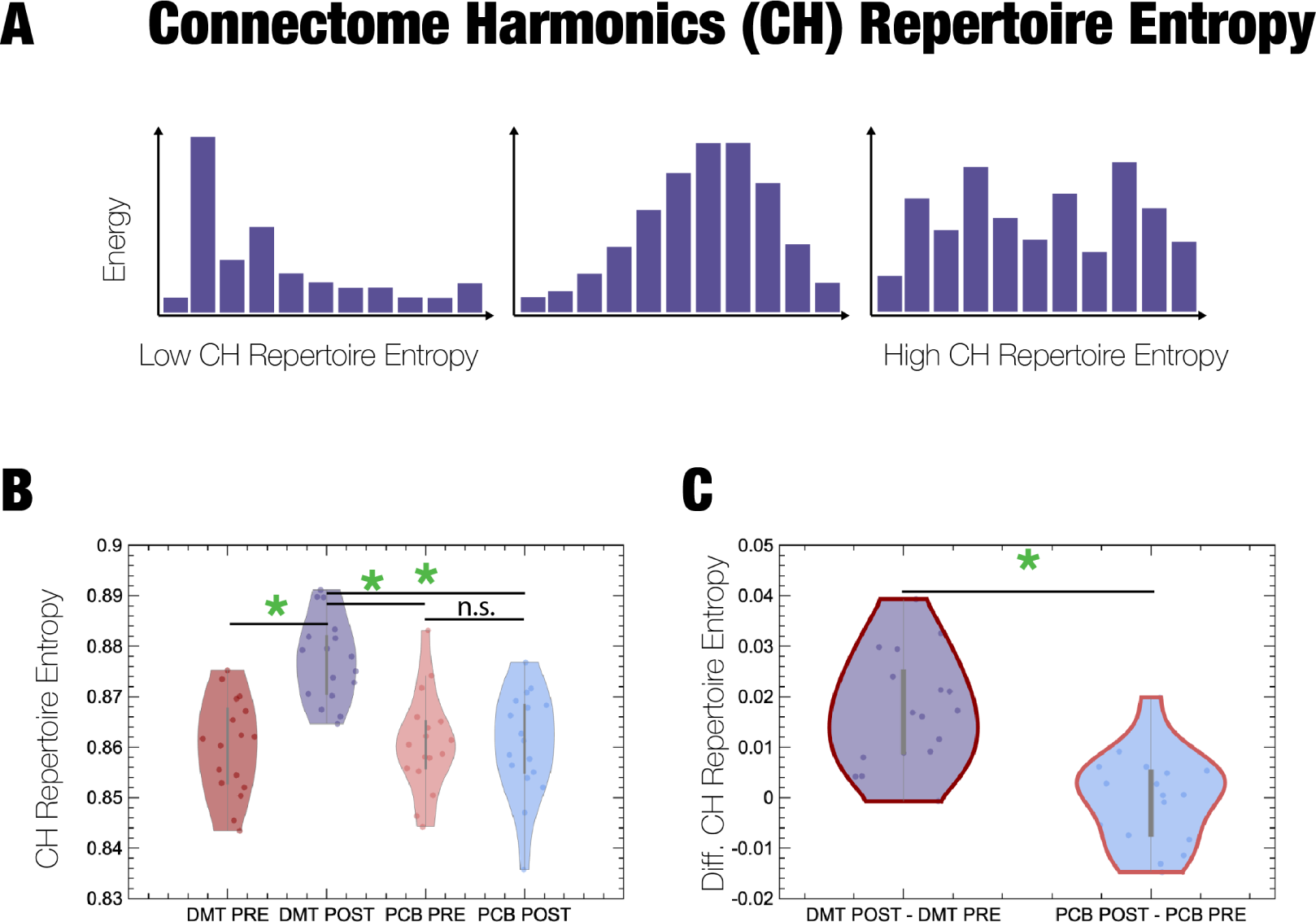
Repertoire of Connectome Harmonics and CH Repertoire Entropy: **A)** Examples of possible repertoires of connectome harmonics. The example on the left has the lowest CH repertoire entropy where the repertoire is dominated by a specific range of spatial frequencies, and the one on the right has the highest entropy where the distribution of the spatial frequencies approaches the uniform distribution with maximum entropy. **B)** CH repertoire Entropy (Pre/Post DMT: p-value < 10^−4^, Pre PCB/Post DMT: p-value < 10^−4^, Post PCB/Post DMT: p-value < 10^−4^ and non-significant difference between Pre/Post PCB, paired t-test). **C)** CH repertoire Entropy Difference (Diff. in Pre-Post DMT and Pre-Post PCB: p-value < 10^−5^, paired t-test).

### Time-resolved coupling of harmonic signatures and subjective experience

Up to this point, we found that DMT replicates the previous observations of increased entropy of the connectome harmonic repertoire under psychedelics. We also found that, like psilocybin and LSD, DMT reduces the contribution of low-frequency harmonics to brain activity, in favour of high-frequency harmonics. Next, we address a further question that these previous studies could not address, but which is made possible by the fast pharmacokinetics and dynamics of IV DMT. Namely, are the changes in connectome harmonic signatures related to changes in subjective experience in a time-resolved manner?

First, we ask whether temporal changes in the entropy of connectome harmonics correlate with temporal changes in the subjective rating of intensity of the DMT experience. We find that this is indeed the case: for half of the individuals, we found significant correlations between the intensity ratings and the DMT session (Spearmann correlation p < 0.05 uncorrected, Figure 5 A and C). Furthermore, on a group level, these individual correlations are statistically significant (ttest p< 0.05). In other words, the changes in repertoire entropy of CH induced by DMT at the neural level, correlate with DMT-induced changes in subjective intensity.

**Fig 5.**
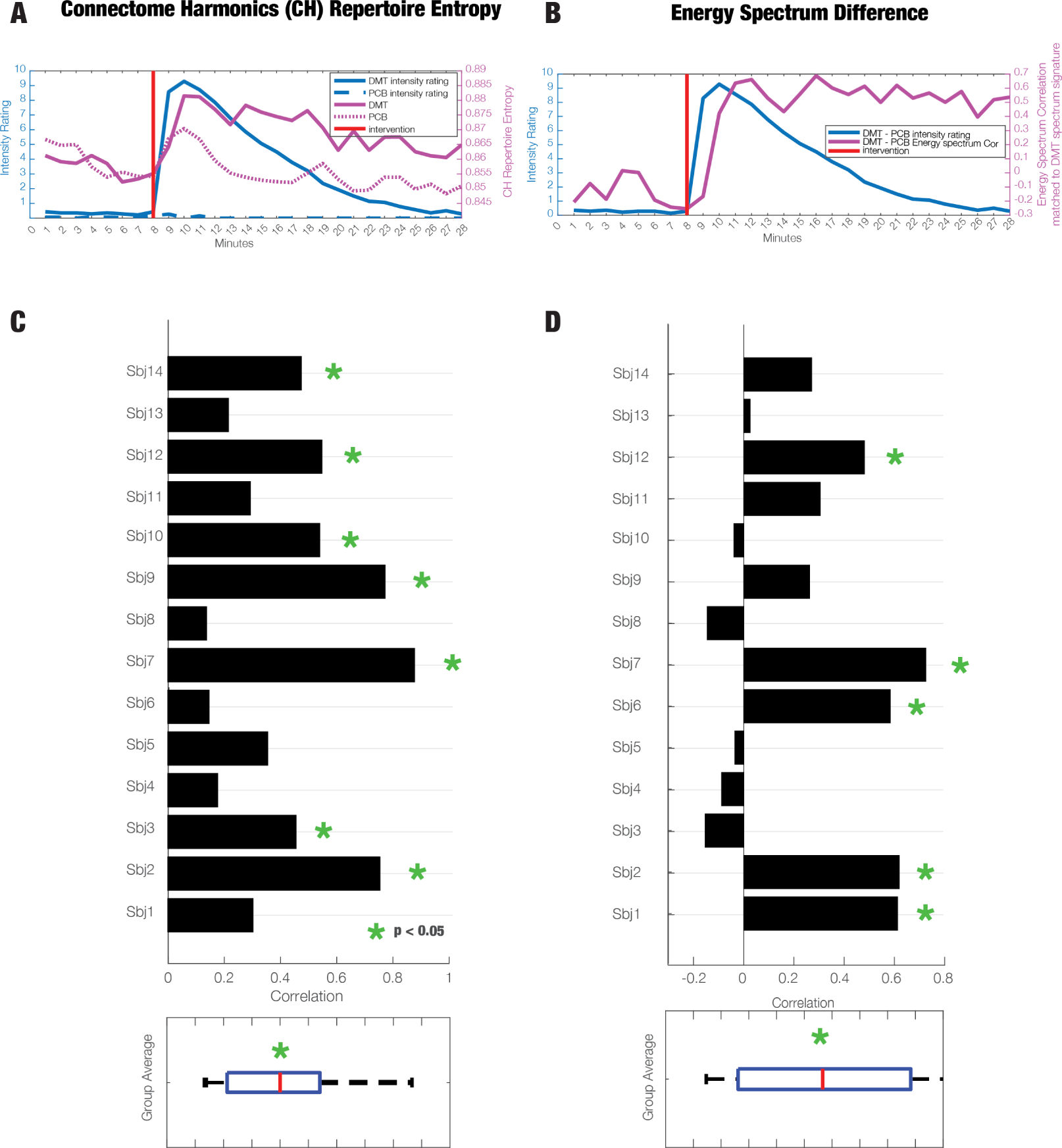
Time-resolved and subject-specific measures of CH Repertoire Entropy and Energy Spectrum Difference in DMT: **A)** The timecourse of CH Repertoire Entropy for the 28 minutes of recording. **B)** The timecourse of Energy Spectrum Difference for the 28 minutes of recording. **C)** Subject specific correlations between the CH Repertoire timecourses of the DMT condition and intensity ratings. The group average of the correlation values between CH Repertoire Entropy and Intensity Ratings is statistical significant (green star, p<0.05). **D)** Subject specific correlations between the Energy Spectrum Difference timecourses and intensity ratings. The group average of the correlation values between CH Repertoire Entropy and Intensity Ratings is statistical significant (green star, p<0.05).

Second, we ask whether the ability to detect the specific connectome harmonic signature of the psychedelic experience (shared by DMT with LSD and psilocybin; Figure 5 B and D) correlates with a more intense subjective experience. Once again, we find that this is the case: for five individuals, the correlations between energy spectrum difference and intensity ratings were significant (Spearmann correlation p < 0.05 uncorrected). Moreover, these correlations were significant at the group level (ttest p< 0.05). In line with CH repertoire entropy, the energy spectrum difference reflects the DMT-induced changes in subjective intensity in a time-resolved manner. We include the time-resolved evolution of the energy spectrum difference in the supplementary information (SI Figure S5).

Traditionally, EEG signatures as described by LZ complexity have been shown to reflect well the DMT-induced subjective intensity in a time-resolved manner.^21^ Here, we wanted to see whether we observe a cross-modal relationship between the different measures of complexity: namely the LZ complexity derived from EEG and CH repertoire entropy from fMRI. We show that indeed it is the case that on the group level the LZ complexity (defined as the difference between DMT and PCB conditions as well DMT alone) correlate significantly with the CH repertoire entropy which is not the case for the placebo condition (Spearmann correlation ** p<0.001, *** p<0.0001 SI Figure S4). However, when comparing “richness of the experience” on a subject level to the magnitude of CH repertoire entropy, we have not observed significant correlation as has been previously shown between “richness of the experience” and LZ complexity^21^ (SI Figure S9).

### Sensitivity and robustness

To ensure the robustness of our results, we replicate our main analysis of DMT CH signature and match with other signatures using 25 logarithmically spaced bins instead of the 15 bins canonically employed for CHD analysis (SI Figure S6).

Conversely, we also show that this ability to replicate results is not merely an indicator that *any* basis function will produce similar results. We illustrate this point by using connectome harmonics obtained from a degree-preserving randomisation of the original structural connectome, which fails to show the loss of energy at low frequencies (SI Figure S8 A), and fails to capture the expected relationship of DMT with ketamine and disorders of consciousness (SI Figure S8 B).

## 1 Discussion

Here, we used connectome harmonic decomposition to represents functional brain signals in terms of their relationship with the detailed network organisation of the human connectome, and understand how this structure-function relationship is altered by the potent psychedelic agent, N,N-Dimethyltryptamine (DMT). Our main hypothesis was that, similar to LSD and psilocybin, DMT would also induce shift away from low to high spatial frequencies of the harmonics. From the connectome harmonic perspective, the spatio-temporal dynamics have been modelled in terms of the temporal contribution of a set of spatial frequencies realised on the structural connectivity of the brain - the connectome. Just like temporal harmonics from traditional Fourier analysis quantify time-dependence in the signal, so the harmonic modes of the human connectome quantify connectome-dependence in brain activity from global to local spatial functional patterns. The results demonstrate full harmonic spectrum changes under the influence of DMT, with a suppression of low-frequency harmonics and an increase of high frequency harmonics. Here, for the first time, fMRI recordings of participants under the influence of the psychedelic N,N-Dimethyltryptamine (DMT) were analysed with this method, revealing time-resolved changes in CH repertoire entropy, thus corroborating previous findings.

The entropic brain hypothesis proposed that the richness of the spatio-temporal dynamics along the entropy continuum,i.e., within a meaningful range, will index the richness of consciousness experience. Furthermore, it proposed^18^ - and later shown^22^ - that the psychedelic-induced state would feature increase in the level of entropy within the brain.^18, 19^ Cognitively, the psychedelic experience has been hypothesised to undermine the integrity of large-scale networks that constrain the susceptibility of any incoming and ‘lower level’ signal; like a wheel constraining the movement of its spokes. The psychedelic state is a state of cognitive and perceptual novelty, as bottom-up signals are put on the same footing with high-level priors, assumptions, or constraining predictive models; in other words, the functional hierarchy is lost under psychedelics, resulting in the so-called ‘anarchic brain’.^23^ Here, we have shown, for the first time, the effect of DMT on repertoire entropy as defined by the connectome harmonic power spectrum. This is supported by an increase in the high-frequency energy spectrum of localised harmonic contributions, and, at the same time, a suppression of the low frequency energy spectrum representative of global harmonic contributions broadly representative of the known large-scale networks.^13^ How structure shapes function has been at the forefront of contemporary neuroscience^24, 25^ with many approaches considered.^26–28^ Recent advances have considered diffusion process to describe the unfolding brain activity on the structural connectome, of which connectome harmonics are the representative example,^13^ but also considered elsewhere^29–31^ as well as on the communication structure.^32, 33^ Furthermore, recent work and an ongoing debate have highlighted the importance of geometry as an important structural feature in shaping the unfolding dynamics.^34–36^ In general terms, the correlation strength of structure-function relationships has been indicative of the level of consciousness - a stronger relationship has signified a loss of consciousnesses, for example in anesthesia.^37–39^ In terms of connectome harmonics one of the potential interpretations has been that low frequency harmonics approximate the global structural topology of the underlying graph, while higher eigenvalues capture localised representations. This is relevant, as the observed effect here is the opposite to the reduced levels of consciousness, with a suppression of lower frequency harmonics and an increase of high frequency harmonics suggesting an opposite trend in reduced levels of consciousness towards a brain state governed by the global (rather than local) organisation of the structural connectome. This leads to the following question: Could the harmonic spectrum be used as a signature of the different brain states of consciousness? This has been explored in a recent study where high generalisibility of the connectome harmonic decomposition spectrum was shown across minimal conscious, anesthetic, and ketamine and LSD-induced psychedelic states.^4^ Meaning, CHD spectrum could be used to categorize these diverse states of consciousness in a predictable and meaningful way.

To represent fMRI activity in different brain states, it is possible to use different bases on which the activity is projected. In this sense, connectome harmonics are considered as an extension of spherical harmonics - similarly derived as the eigenfunctions of the Laplace operator applied to the sphere.^34, 40, 41^ Hence, when considering only local grey-matter connectivity, connectome harmonics equal to spherical harmonics represented on the cortical surface. However, we argue here that long-range connectivity is a necessary feature for an accurate representation of brain states.^42, 43^ Therefore, connectome harmonics are extending spherical harmonics approaches by embedding both local grey matter and long-range white matter connectivity of the human brain. Moreover, when the underlying graph is randomised (even as the number of connections of each node is preserved), the ability to correctly identify brain states corresponding to the psychedelic state versus loss of consciousness is lost (SI Figure S8) consistently with what was previously observed^4^.

Experimentally, the DMT dataset is a single-blind and counter-control placebo design and, unlike previous experiments with psilocybin^10^ and LSD,^11^ contains a control group that is important to differentiate the changes in the connectome harmonic decomposition under the influence of DMT from its baseline. Moving forward, future work might further differentiate the level of vigilance that comes with the psychedelic experience by considering additional control groups under the influence of stimulants such as Modafinil and caffeine. Recently, this was done with methylphenidate, controlling for arousal. Methylphenidate matched psilocybin in its pro-arousal effects but failed to show the marked characteristic brain function changes we have become accustomed to with the classic psychedelics.^44^

## 2 Summary

To summarise, we have analysed the DMT-induced state, through the lens of Connectome Harmonic Decomposition. Importantly, this description allows to represent the spatio-temporal dynamics and its associated psychedelic-induced state as a time-varying contribution of fundamental building blocks of brain activity: different scales of the structural connectome. The results show that a suppression of low-frequency harmonics and an increase of high frequency harmonics is observed in the DMT-induced state, consistent with the effects of other psychedelics, and corresponding to a decoupling of function from the underlying structure.

Notably, leveraging the rapid and well-characterised temporal sequence of DMT subjective effects, we were also able to demonstrate that the time-resolved changes in CHD signature induced by LSD, are predictive of time-resolved changes in subjective intensity of the psychedelic experience. Likewise, DMT induced an increase int he entropy of the connectome harmonic repertoire, which was predictive of time-resolved changes in subjective intensity. These time-resolved results demonstrate with unprecedented resolution the close relationship between connectome harmonics and subjective experience - linking brain and mind.

## 3 Supplementary Material

**Fig S1.**
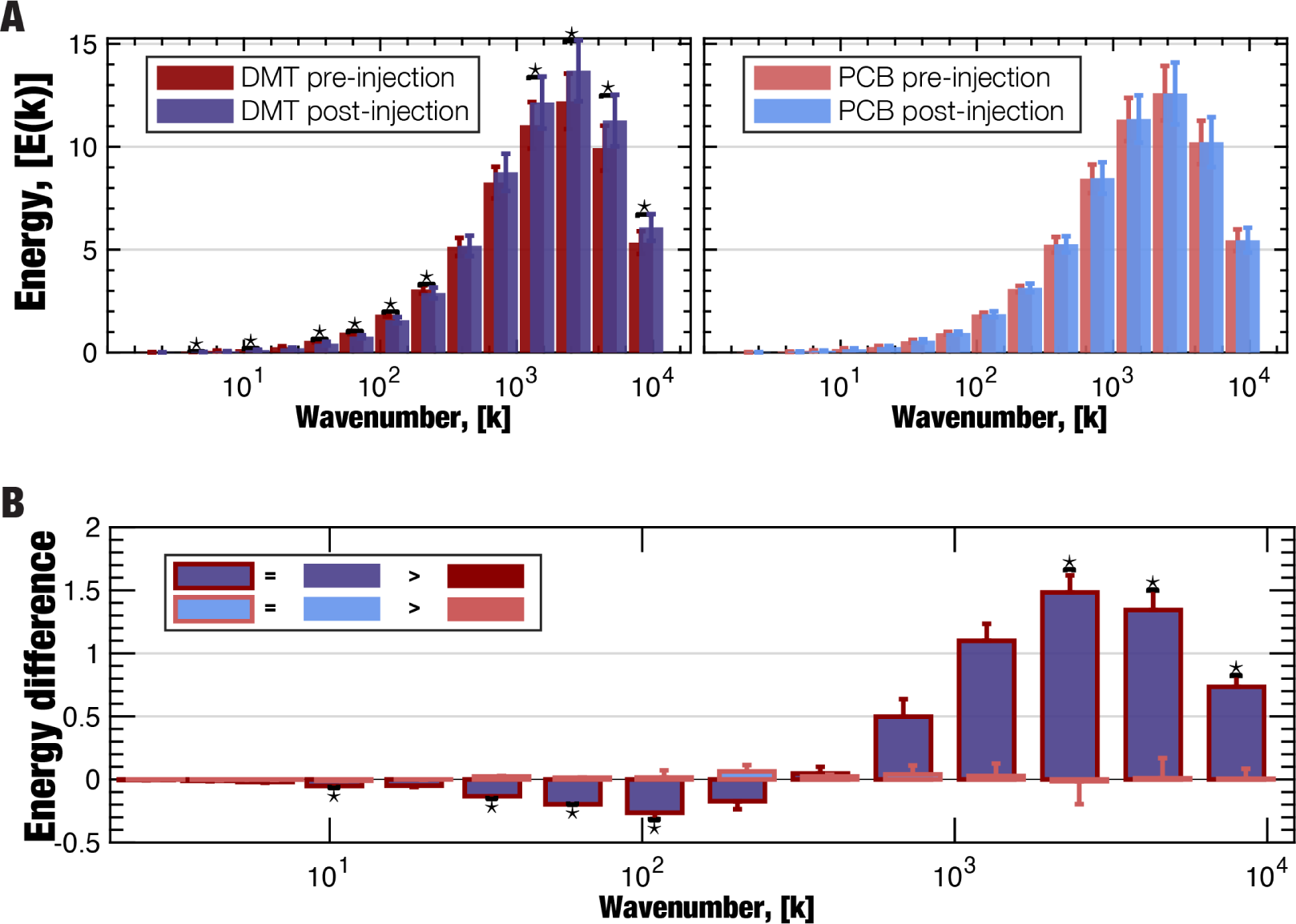
**A** Replication of DMT CH energy signature using dense structural connectome from 985 HCP participant. **B** Replication of DMT CH energy difference signature using dense structural connectome from 985 HCP participant.

**Fig S2.**
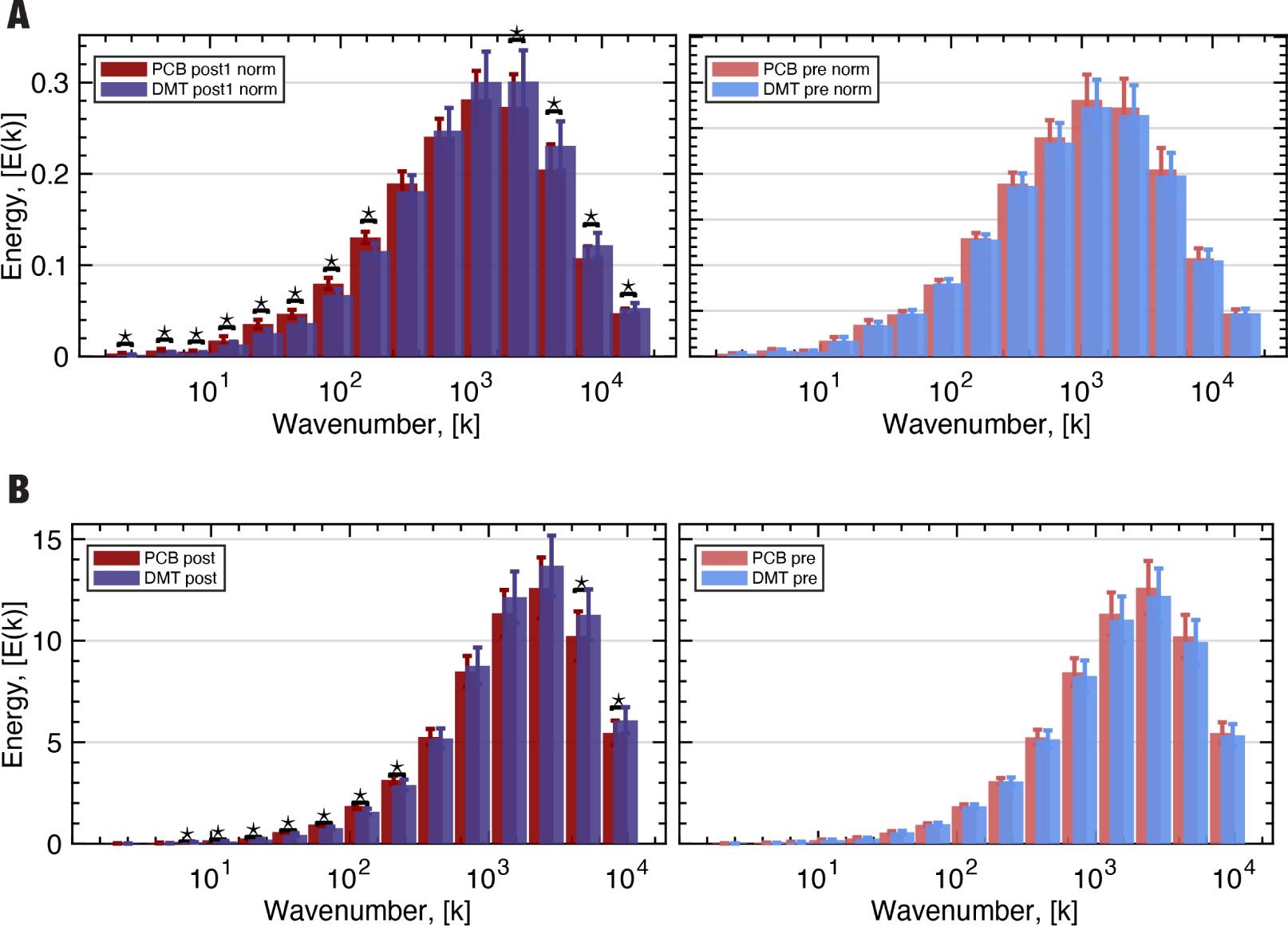
**A** The comparison of post DMT conditions of CH energy signature using the original connectome. **B** Replication of the post DMT condition of CH energy signature using dense structural connectome from 985 HCP participant.

**Fig S3.**
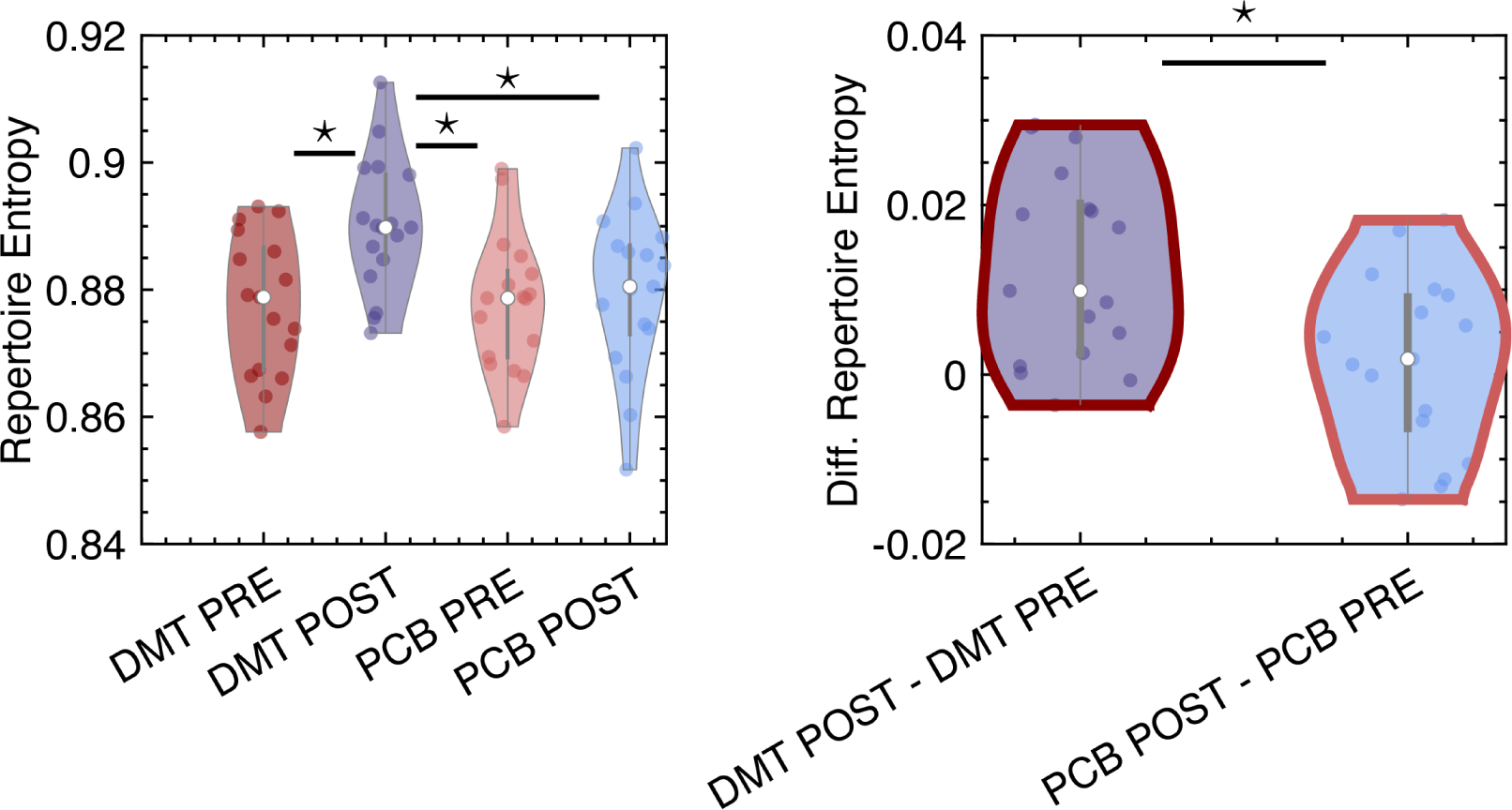
**A** Replication of DMT CH repertoire entropy using dense structural connectome from 985 HCP participant. Repertoire Entropy (Pre/Post DMT: p-value < 10^−4^, Pre PCB/Post DMT: p-value < 10^−4^, Post PCB/Post DMT: p-value < 10^−4^ and non-significant difference between Pre/Post PCB, paired t-test). Repertoire Entropy Difference (Diff. in Pre-Post DMT and Pre-Post PCB: p-value < 10^−5^, paired t-test).

**Fig S4.**
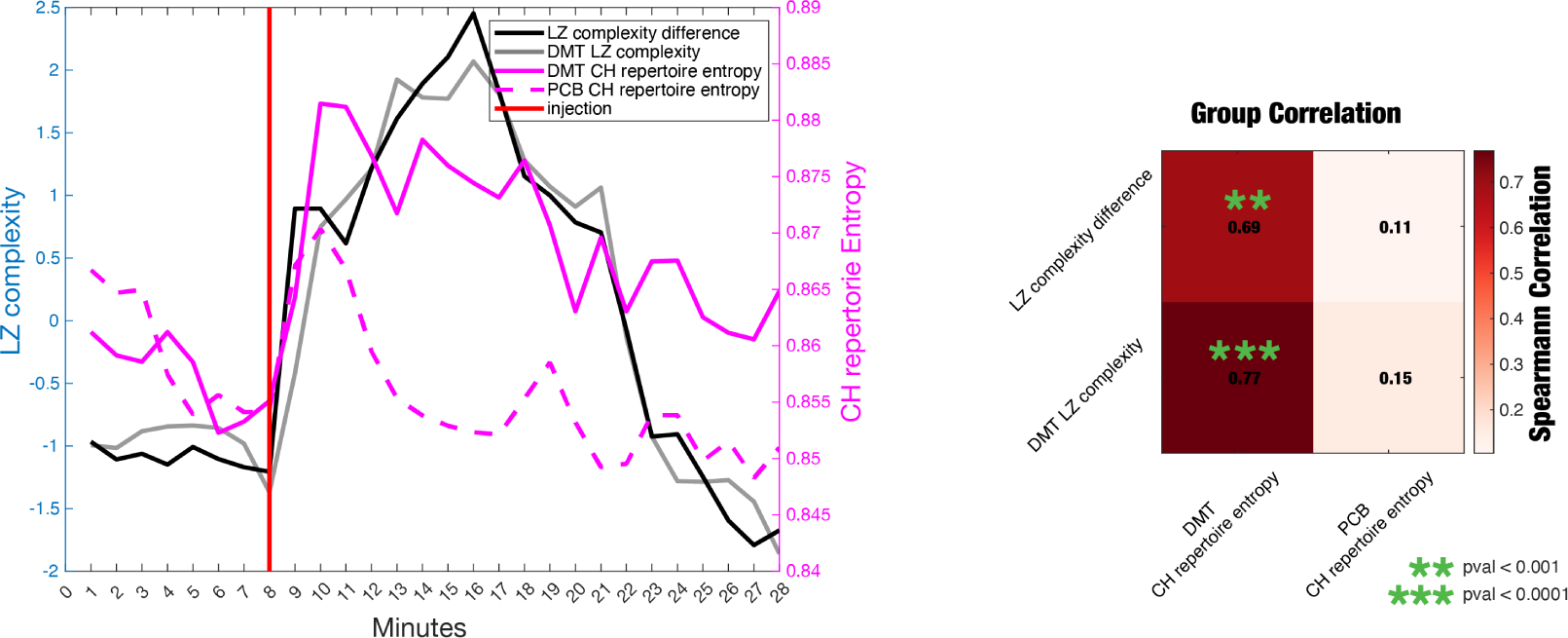
FMRI-based CH repertoire entropy correlates with EEG-based LZ complexity. Both the subject-averaged DMT LZ complexity and DMT-PCB complexity biomarkers from^21^ significantly correlate with the CH repertoire entropy measures of this study (LZ complexity difference vs. DMT CH repertoire entropy p-val <0.001, DMT LZ complexity vs. DMT CH repertoire entropy p-val <0.0001)

**Fig S5.**
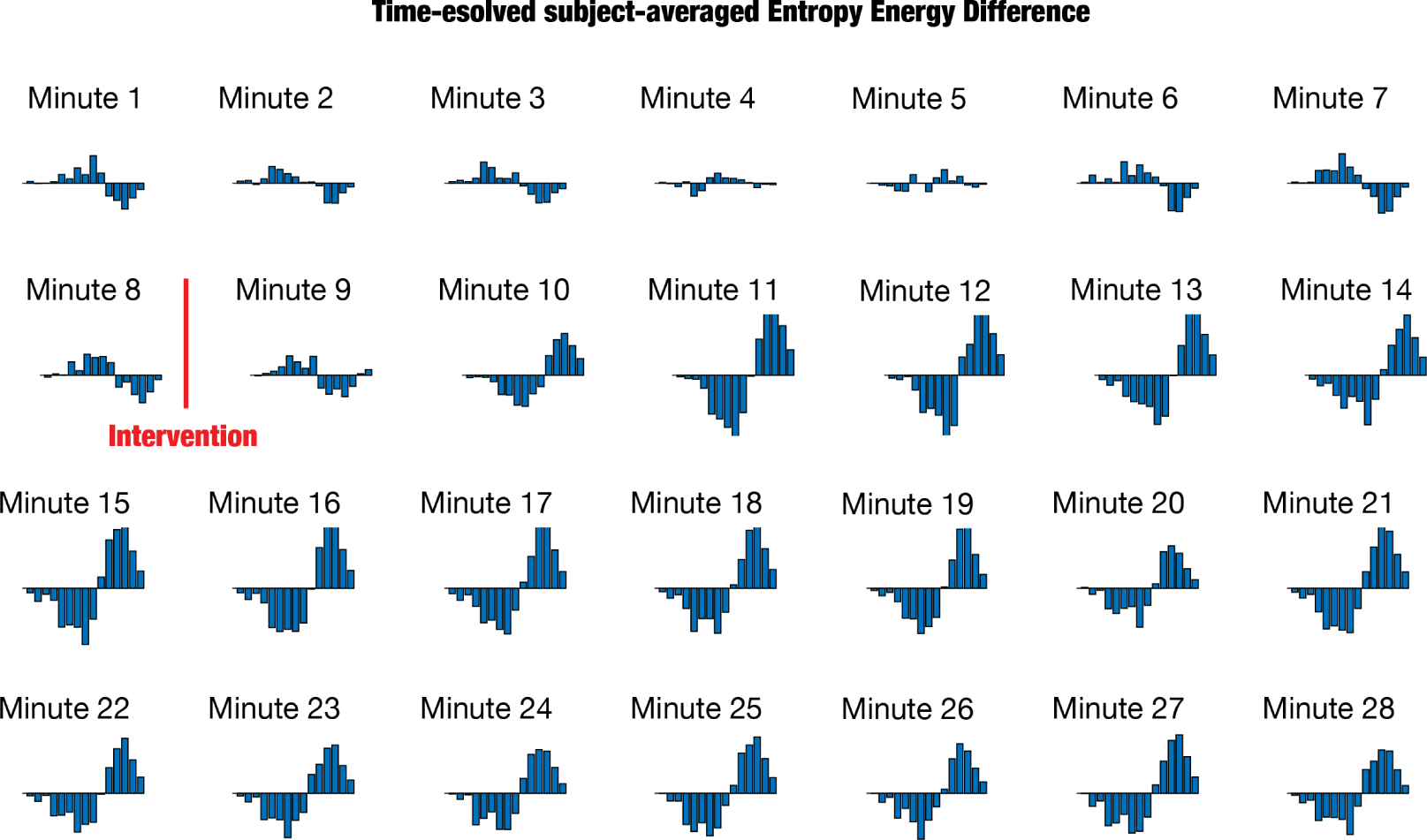
Time-unfolded Energy Spectrum Difference. Time-resolved measure of Energy Spectrum Difference across the 28 minutes of recording averaged across all the subjects

**Fig S6.**
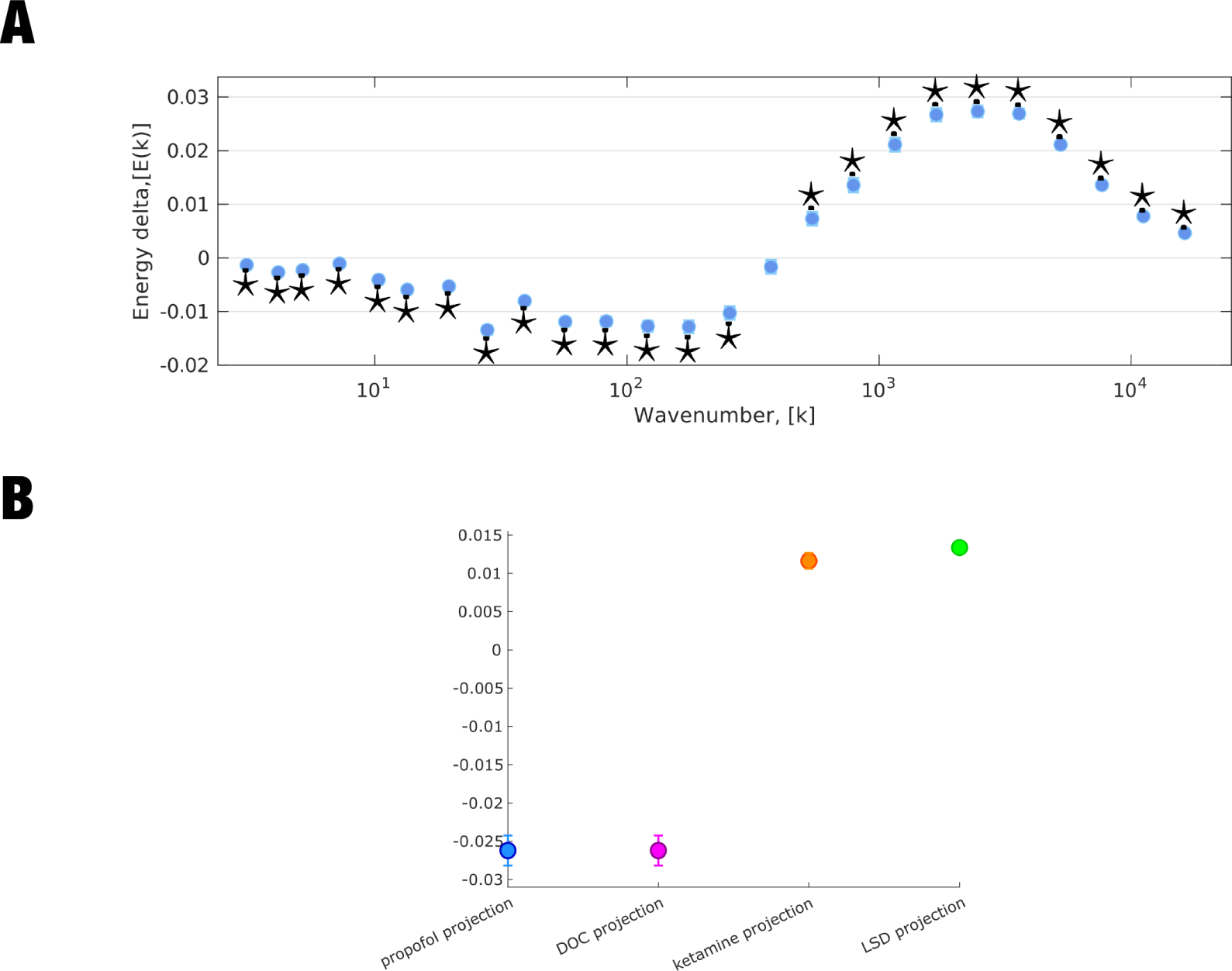
**A** Replication of DMT CH energy signature using 25 bins. **B** Replication of the relationship between multivariate CH signatures of DMT and other states of altered consciousness, using 25 bins. Plots show the fixed effects (and 95% CI) of projections (dot product) between the multivariate connectome harmonic signature of DMT, and four other states previously investigated by:^4^ anaesthesia (blue), DOC patients (violet), ketamine (orange), and LSD (green).

**Fig S7.**
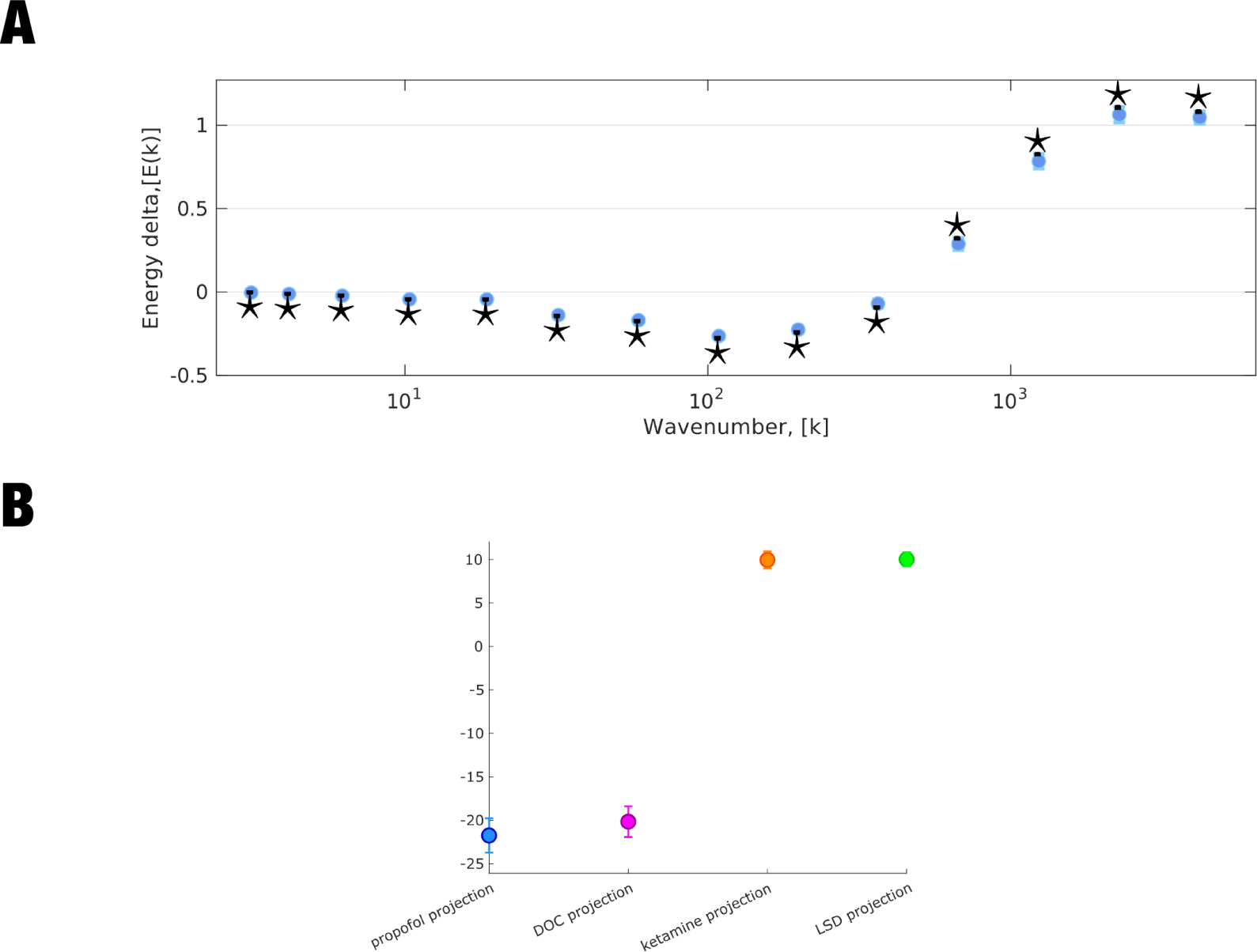
**A** Replication of DMT CH energy signature using a high-resolution human connectome from N=985 healthy individuals. **B** Replication of the relationship between multivariate CH signatures of DMT and other states of altered consciousness, using a high-resolution human connectome from N=985 healthy individuals. Plots show the fixed effects (and 95% CI) of projections (dot product) between the multivariate connectome harmonic signature of DMT, and four other states previously investigated by:^4^ anaesthesia (blue), DOC patients (violet), ketamine (orange), and LSD (green).

**Fig S8.**
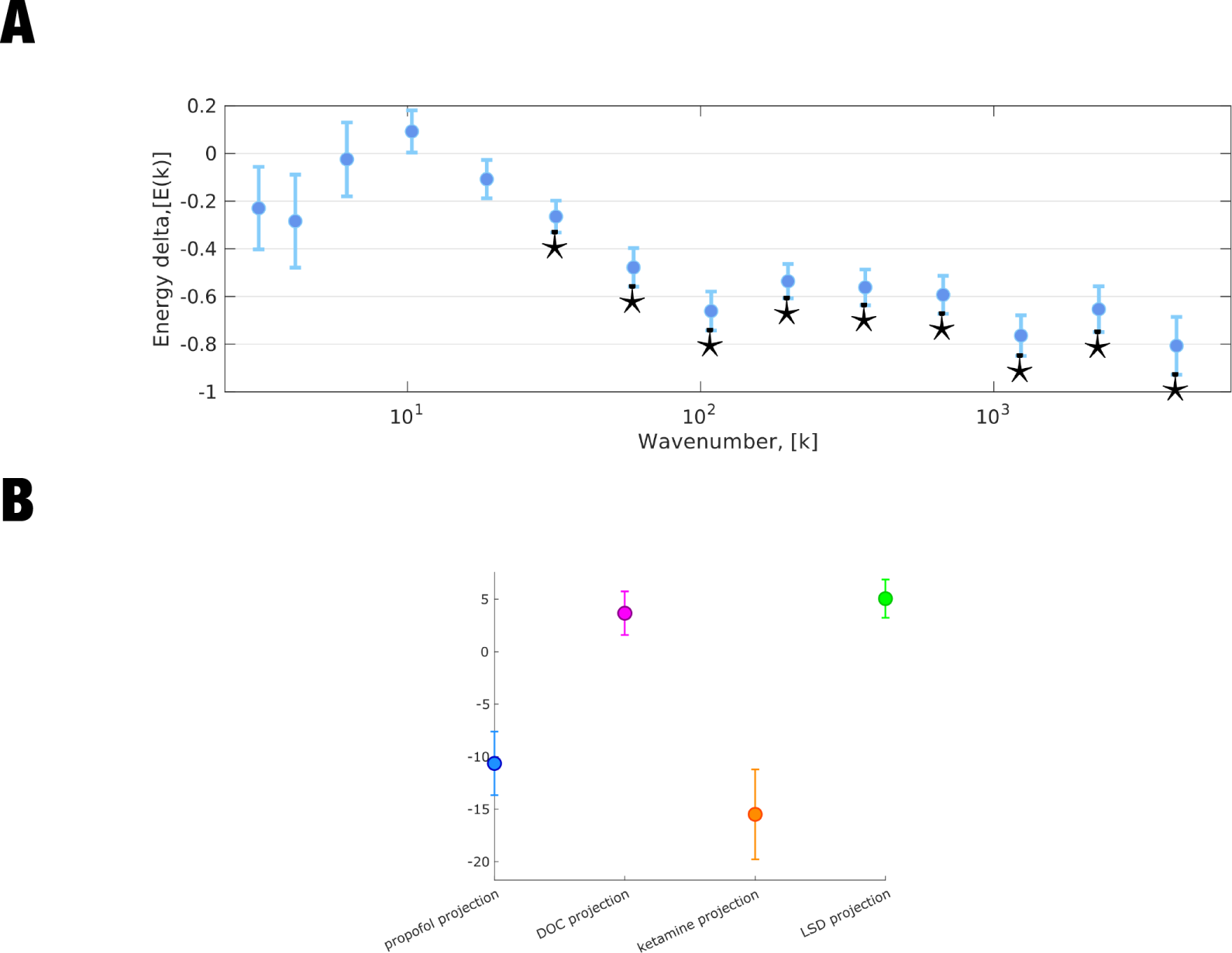
**A** Replication of DMT CH energy signature using connectome harmonics from a degree-preserving randomised connectome. **B** Replication of the relationship between multivariate CH signatures of DMT and other states of altered consciousness, using connectome harmonics from a degree-preserving randomised connectome. Plots show the fixed effects (and 95% CI) of projections (dot product) between the multivariate connectome harmonic signature of DMT, and four other states previously investigated by:^4^ anaesthesia (blue), DOC patients (violet), ketamine (orange), and LSD (green).

**Fig S9.**
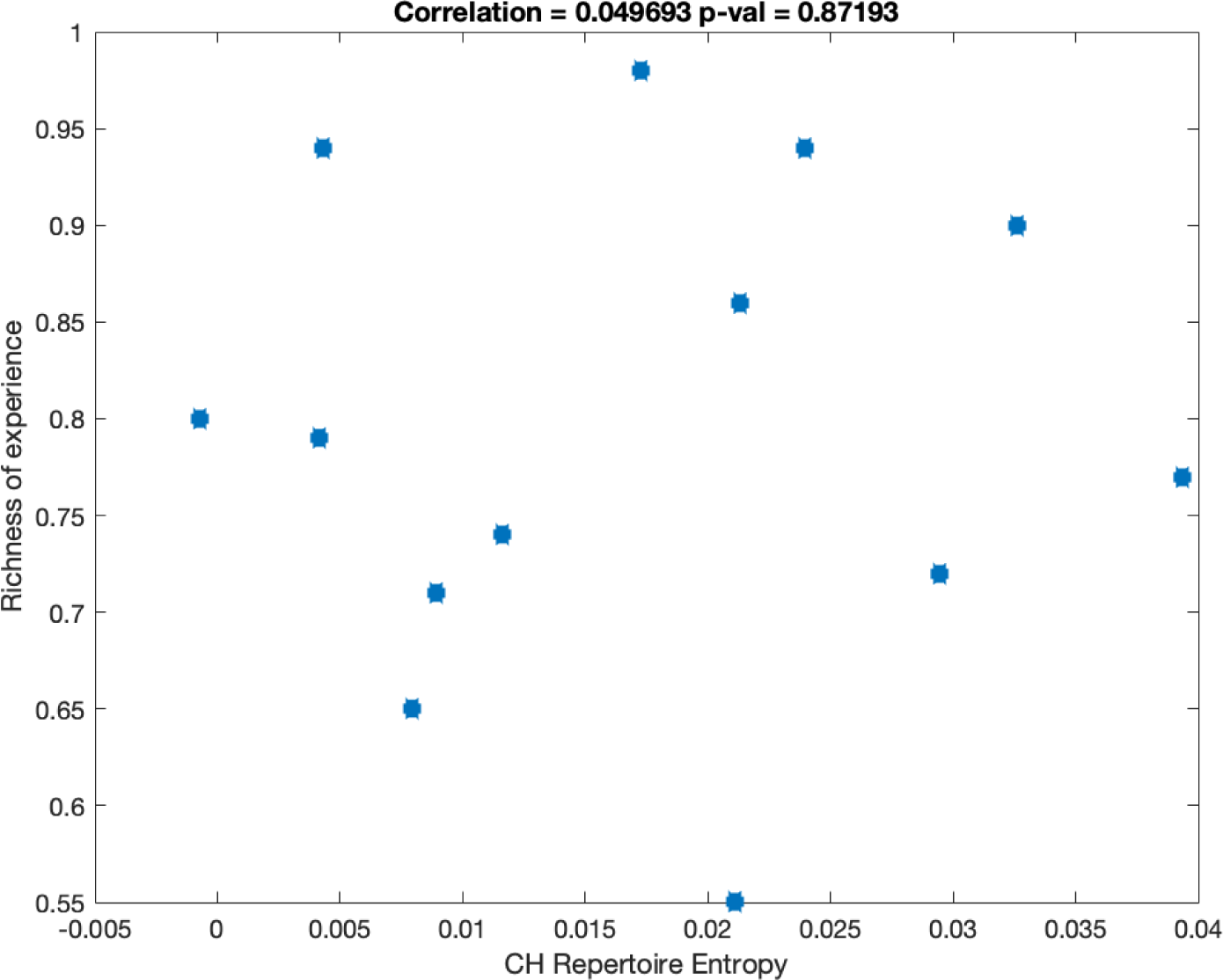
**A** Correlation between CH Repertoire Entropy and “richness of the experience” under the DMT-induced state. Unlike in^21^ where LZ complexity has been associated with “richness of the experience”, here we report a non-significant Spearmann correlation between CH Repertoire Entropy and “richness of the experience” (corr = 0.05, p-val = 0.87).

